## Supplementary Information for "Vancomycin-resistant enterococci colonise the antibiotic-treated intestine by occupying overlapping but distinct nutrient- and metabolite-defined intestinal niches"

### SUPPLEMENTARY MATERIAL:

### SUPPLEMENTARY TABLES

**Table S1: VRE strains were resistant to broad-spectrum antibiotics that promote VRE intestinal colonisation.** The broth microdilution technique was used to measure the MICs of *E. faecalis* and *E. faecium* for vancomycin (VAN), metronidazole (MTZ), ceftriaxone (CRO), piperacillin/tazobactam (TZP) and clindamycin (CLI). Assays were performed under aerobic conditions in Mueller-Hinton broth following EUCAST standards (n = 3).

| Strain | VAN (mg/L) | MTZ (mg/L) | CRO (mg/L) | TZP (mg/L) | CLI (mg/L) |
|---|---|---|---|---|---|
| *E. faecalis* (NCTC 12201) | > 128 (resistant) | 32 (resistant) | > 128 (resistant) | 8 (resistant) | > 128 (resistant) |
| *E. faecium* (NCTC 12202) | 8 (resistant) | 32 (resistant) | 8 (resistant) | 64 (resistant) | > 128 (resistant) |

**Table S2: Comparison of *E. faecium* (NCTC 12202) or *E. faecalis* (NCTC 12201) growth in faecal cultures treated with different antibiotics (data from Figs. 1c and 1d).** Mixed effects model of log transformed CFU/ml with Tukey's multiple comparisons test. * = P ≤ 0.05, ** = P ≤ 0.01, *** = P ≤ 0.001, **** = P ≤ 0.0001.

| Comparison | *E. faecium* | | *E. faecalis* | |
|---|---|---|---|---|
| | Adjusted P value | Summary | Adjusted P value | Summary |
| Faeces + MTZ vs. Faeces + CLI | 0.0858 | ns | 0.0008 | *** |
| Faeces + MTZ vs. Faeces + VAN | 0.0129 | * | <0.0001 | **** |
| Faeces + MTZ vs. Faeces + CRO | <0.0001 | **** | <0.0001 | **** |
| Faeces + CLI vs. Faeces + VAN | 0.6969 | ns | 0.4509 | ns |
| Faeces + CLI vs. Faeces + CRO | 0.0008 | *** | <0.0001 | **** |
| Faeces + VAN vs. Faeces + CRO | 0.0038 | ** | 0.0009 | *** |

**Table S3: Half maximal inhibitory concentrations ($IC_{50}$) for microbial metabolites and VRE growth.** $IC_{50}$ values indicate the concentration of the metabolite that is required to inhibit 50% of vancomycin-resistant *E. faecium* or *E. faecalis* growth. N/A = not available or not applicable.

| Metabolite | *E. faecium* | | | *E. faecalis* | | |
|---|---|---|---|---|---|---|
| | NCTC 12202 | NCTC 12204 | DSM 25698 | NCTC 12201 | NCTC 13779 | DSM 116626 |
| Formate | 49.2 | 47.5 | 27.8 | 117.8 | 36.9 | 41.5 |
| Acetate | 188.7 | 129.3 | 77.6 | 266.9 | 143.5 | 131.8 |
| Propionate | 66.7 | 49.6 | 26.5 | 36.4 | 20.7 | 40.2 |
| Butyrate | 48.0 | 65.0 | 31.7 | 80.0 | 38.4 | 69.4 |
| Valerate | 13.9 | 23.8 | 7.7 | 19.1 | 18.4 | 31.7 |
| Isobutyrate | 155.5 | 67.1 | 32.4 | 160.4 | 21.1 | 132.8 |
| Isovalerate | 50.7 | 19.4 | 15.5 | 56.9 | 17.5 | 36.8 |
| Lactate | 138.8 | 18.3 | 119.5 | NA | 134.6 | NA |
| 5-aminovalerate | > 512 | N/A | N/A | N/A | N/A | N/A |
| Ethanol | N/A | 119.3 | > 512 | N/A | 134.7 | N/A |

**Table S4:** Reagents and oligonucleotides used in this study.

| Reagent | Company | Code |
|---|---|---|
| 3-Trimethylsilylpropionic-2,2,3,3-$d_4$ acid sodium salt | Merck | 269913-1G |
| 5-aminovalerate | Merck | 123188-5G |
| Acetic acid | Merck | A6283-100ML |
| Adenosine | TCI Europe | A0152-5G |
| Agar | VWR | 84609.05 |
| Ammonium sulfate | Sigma | A4418-500G |
| Arabinogalactan | TCI Europe | A1328-100G |
| Bile salts | Fisher Scientific | 10128872 |
| Boric acid | Sigma | B6768-1KG |
| Brain heart infusion broth | Sigma | 53286-500G |
| Brilliance CRE agar | Fisher Scientific | 13205349 |
| Brilliance VRE agar | Fisher Scientific | 11924112 |
| Butyric acid | Merck | B103500-100ML |
| Calcium chloride anhydrous | Fisher Scientific | 10515671 |
| Casein | VWR | A13707.36 |
| Casein hydrolysate | Merck | 22090-100G |
| Cobalt (II) chloride | Sigma | C8661-25G |
| Columbia Agar | VWR | 84621.05 |
| Copper (II) chloride | Sigma | C3279-100G |
| Defibrinated horse blood | SLS | MED1306 |
| Deuterium oxide | Goss Scientific | DLM-4-100G |
| D-Fructose | Merck | F0127-100G |
| D-Galactose | Fluorochem | 218380-25G |
| D-Glucose | VWR | 101174Y |
| D-Maltose | TCI Europe | M0037-25G |
| D-Mannose | TCI Europe | M0037-25G |
| DNeasy PowerLyzer PowerSoil Kit | Qiagen | 12855-100 |
| D-Ribose | Fluorochem | 078886-25G |
| D-Trehalose dihydrate | Fluorochem | 243212-25G |
| D-Xylose | Fluorochem | 209047-100G |
| *Escherichia coli* DNA | Merck | D4889-1UN |
| Ethanol | Fisher Scientific | 10437341 |
| Ethylenediaminetetraacetic acid | Sigma | E5134-50G |
| Fastidious anaerobic agar | SLS | MED1332 |
| Formic acid | VWR | 20320.295 |
| Gifu anaerobic broth | VWR | HIMEM1801-500G |
| Guanosine | TCI Europe | G0171-5G |
| Hemin | Merck | 51280-1G |
| Hydrochloric acid | VWR | 20252.29 |
| Inulin | VWR | A18425.18 |
| Iron sulphate | Sigma | 7782-63-0 |
| Isobutyric acid | Merck | I1754-100ML |
| Isovaleric acid | Merck | 129542-100ML |

| Reagent | Company | Code |
|---|---|---|
| Kao and Michayluk vitamin solution | Merck | K3129 |
| L-Alanine | TCI Europe | A0179-25G |
| L-Arabinose | Fluorochem | 078969-25G |
| L-Arginine hydrochloride | TCI Europe | A0528-25G |
| L-Aspartic acid | TCI Europe | A0546-25G |
| L-Cysteine hydrochloride monohydrate | Merck | C7880-100G |
| L-Fucose | Fluorochem | 216288-5G |
| L-Glutamic acid | TCI Europe | G0059-25G |
| L-Glycine | TCI Europe | G0099-25G |
| L-Isoleucine | TCI Europe | I0181-25G |
| L-Leucine | TCI Europe | L0029-25G |
| L-Lysine hydrochloride | TCI Europe | L0071-25G |
| L-Methionine | TCI Europe | M0099-25G |
| L-Phenylalanine | TCI Europe | P0134-25G |
| L-Proline | TCI Europe | P0481-25G |
| L-Threonine | TCI Europe | T0230-25G |
| L-Tryptophan | TCI Europe | T0541-25G |
| L-Tyrosine | TCI Europe | T0550-25G |
| L-Valine | TCI Europe | V0014-25G |
| Magnesium chloride | Sigma | 7786-30-3 |
| Magnesium sulphate heptahydrate | Fisher Scientific | 10424511 |
| Manganese (II) chloride | Sigma | M5005-100G |
| Menadione | Merck | M9429-25G |
| MiSeq Reagent Kit V3 | Illumina | MS-102-3003 |
| Mucin from porcine stomach type II | Merck | M2378-500G |
| Muller-Hinton broth | Merck | 70192-100G |
| N-acetylglucosamine | TCI Europe | A0092-25G |
| NEBNext Library Quant Kit for Illumina | New England Biolabs | E7630L |
| Nickel dichloride | Sigma | 654507-5G |
| Nicotinic acid | Sigma | 72309-100G |
| Pantethine | Sigma | P2126-1G |
| PCR grade water | Merck | 3315932001 |
| Pectin citrus | VWR | J61021.36 |
| Peptone water | Merck | 70179-500G |
| Platinum Supermix with ROX | Life Technologies | 11730017 |
| Potassium phosphate dibasic | Merck | P3786-500G |
| Potassium phosphate monobasic | Merck | P5655-500G |
| Propionic acid | Merck | P1386-500ML |
| Proteose peptone | Merck | 82450-100G |
| RMPI-1640 amino acid solution | Merck | R7131 |
| SequalPrep Normalisation Plate Kit | Life Technologies | A1051001 |
| Sodium azide | Merck | 71289-5G |
| Sodium bicarbonate | Merck | S6014-500G |
| Sodium chloride | Merck | S7653-1KG |
| Sodium DL-lactate | Merck | 71720-5G |

| Reagent | Company | Code |
|---|---|---|
| Sodium fumerate | Fluorochem | F862900-25G |
| Sodium hydroxide | Fisher Scientific | 10478030 |
| Sodium phosphate dibasic | Merck | 71640-250G |
| Sodium phosphate monobasic | Fisher Scientific | 389872500 |
| Sodium pyruvate | SLS | P2256-5G |
| Sodium succinate dibasic hexahydrate | SLS | S2378-100G |
| Soluble starch | SLS | CHE3622 |
| Starch | Merck | S5127-5KG |
| Sucrose | Merck | 84100-250G |
| Thymine | TCI Europe | T0234-5G |
| Tri-sodium citrate monobasic | VWR | DIUKTR19204-535 |
| Tryptic Soy Broth | Sigma | 22092-500G |
| Tween 80 | Sigma | P1754-25ML |
| Uracil | TCI Europe | U0013-25G |
| Valeric acid | Merck | 240370-100ML |
| Xylan | Universal Biologicals | 38500.02 |
| Yeast extract | Merck | 70161-500G |
| Zinc chloride | Sigma | Z0152-100G |
| 16S sequencing primer: 28F-YM (forward primer, MiSeq adapter sequences in bold) | Eurofins | 5'-**TCGTCGGCAGCGTCAGATGTGTATAAGAGACAG**GAGTTTGATYMTGGCTCAG-3' |
| 16S sequencing primer: 28F-Borrellia (forward primer, MiSeq adapter sequences in bold) | Eurofins | 5'-**TCGTCGGCAGCGTCAGATGTGTATAAGAGACAG**GAGTTTGATCCTGGCTTAG-3' |
| 16S sequencing primer: 28F-Chloroflex (forward primer, MiSeq adapter sequences in bold) | Eurofins | 5'-**TCGTCGGCAGCGTCAGATGTGTATAAGAGACAG**GAATTTGATCTTGGTTCAG-3' |
| 16S sequencing primer: 28F-Bifdo (forward primer, MiSeq adapter sequences in bold) | Eurofins | 5'-**TCGTCGGCAGCGTCAGATGTGTATAAGAGACAG**GGGTTCGATTCTGGCTCAG-3' |
| 16S sequencing primer: 388R (reverse primer, MiSeq adapter sequences in bold) | Eurofins | 5'-**GTCTCGTGGGCTCGGAGATGTGTATAAGAGACAG**TGCTGCCTCCCGTAGGAGT-3' |
| 16S rRNA gene qPCR: BactQUANT forward primer | Eurofins | 5′-CCTACGGGAGGCAGCA-3′ |
| 16S rRNA gene qPCR: BactQUANT reverse primer | Eurofins | 5′-GGACTACCGGGTATCTAATC-3′ |
| 16S rRNA gene qPCR: BactQUANT probe | Life Technologies | 4316034, [6FAM] 5′-CAGCAGCCGCGGTA-3′ [MGBNFQ] |

**Table S5:** Concentrations of microbial metabolites tested in the metabolite inhibition assays. Concentrations were based on measurements taken from faecal samples from 12 healthy human donors from Yip *et al.* (2023)[1].

| Metabolite | Lowest concentration (mM) | Average concentration (mM) | Highest concentration (mM) |
|---|---|---|---|
| Formate | 0.05 | 0.10 | 0.15 |
| Acetate | 10 | 65 | 120 |
| Propionate | 5 | 15 | 35 |
| Butyrate | 3 | 15 | 40 |
| Valerate | 0.5 | 3.5 | 12 |
| Isobutyrate | 0.5 | 2 | 5 |
| Isovalerate | 0.5 | 2 | 5 |
| Lactate | 0.3 | 0.7 | 1.2 |
| 5-aminovalerate | 0.3 | 1 | 4 |
| Ethanol | 0.6 | 10 | 60 |

### SUPPLEMENTARY FIGURES

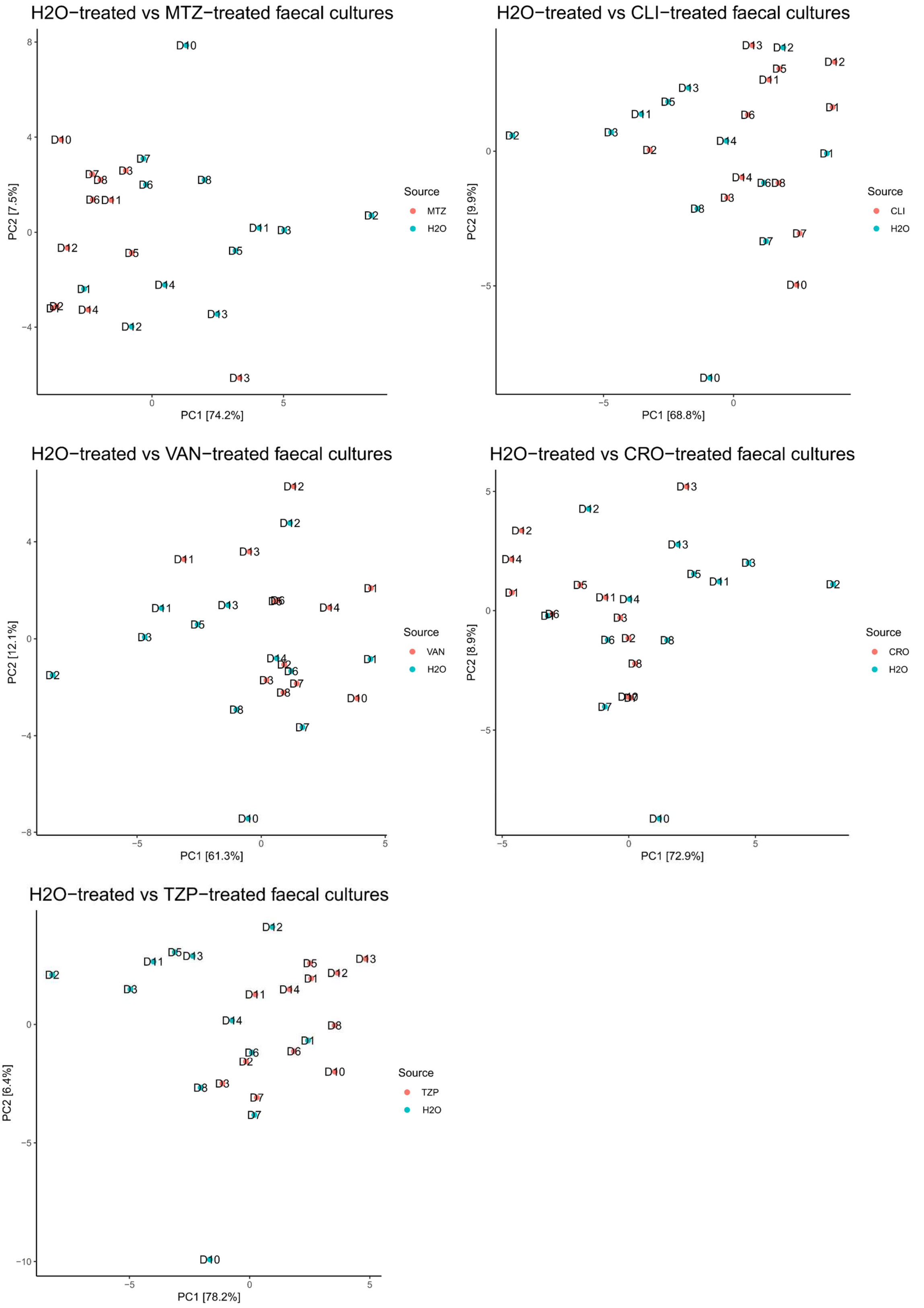


**<u>Figure S1</u>: Principal component analysis (PCA) plots of water-treated vs antibiotic-treated faecal culture samples.** Plots used log-transformed absolute abundance data at the family level. MTZ = metronidazole, CLI = clindamycin, VAN = vancomycin, CRO = ceftriaxone, TZP = piperacillin/tazobactam. D1 - D12 = Donor 1 – Donor 12.

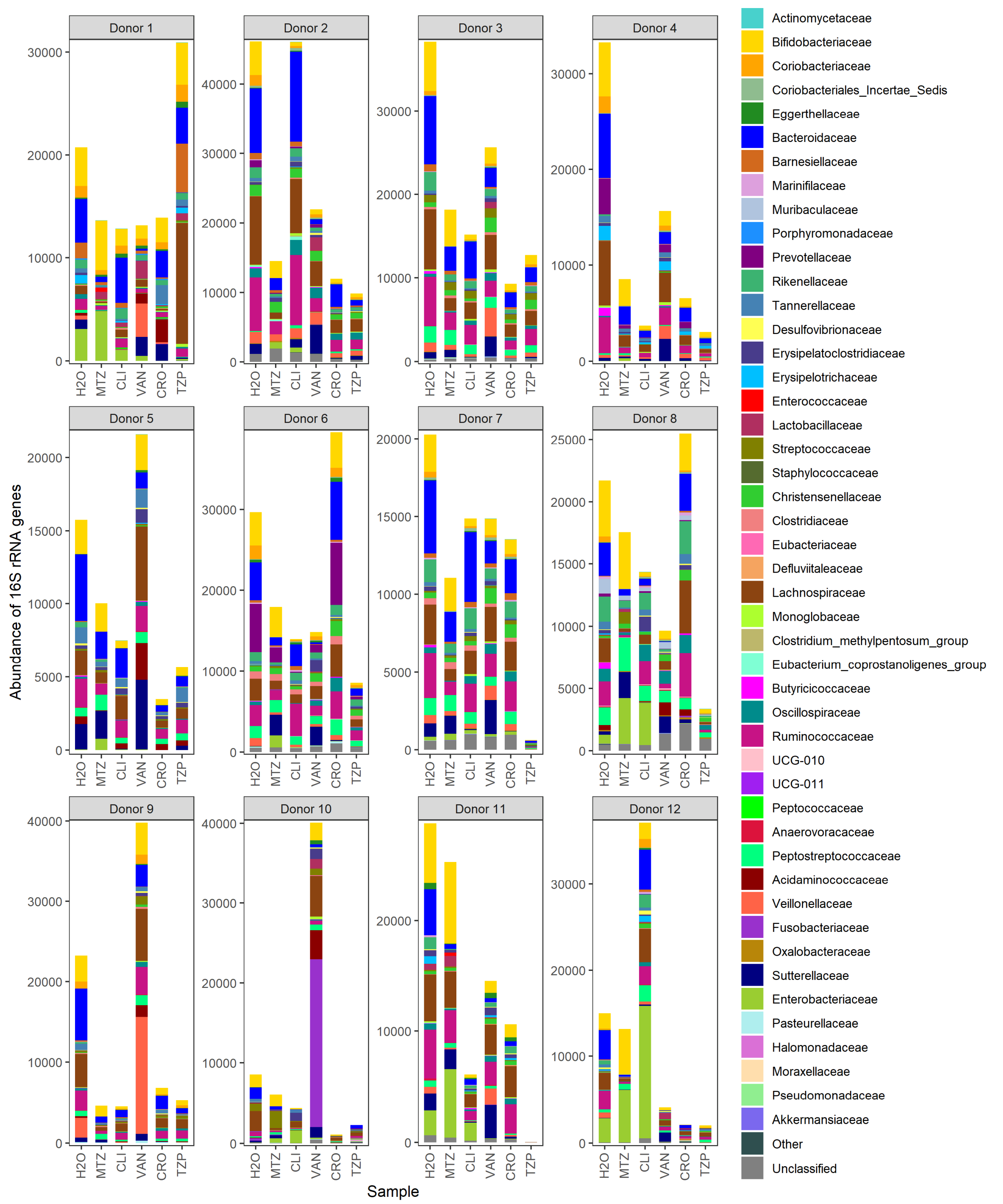


**<u>Figure S2</u>: Stacked bar plots showing the absolute abundances of bacterial families found in water-treated and antibiotic-treated faecal cultures, presented by individual donors.** MTZ = metronidazole, CLI = clindamycin, VAN = vancomycin, CRO = ceftriaxone, TZP = piperacillin/tazobactam.

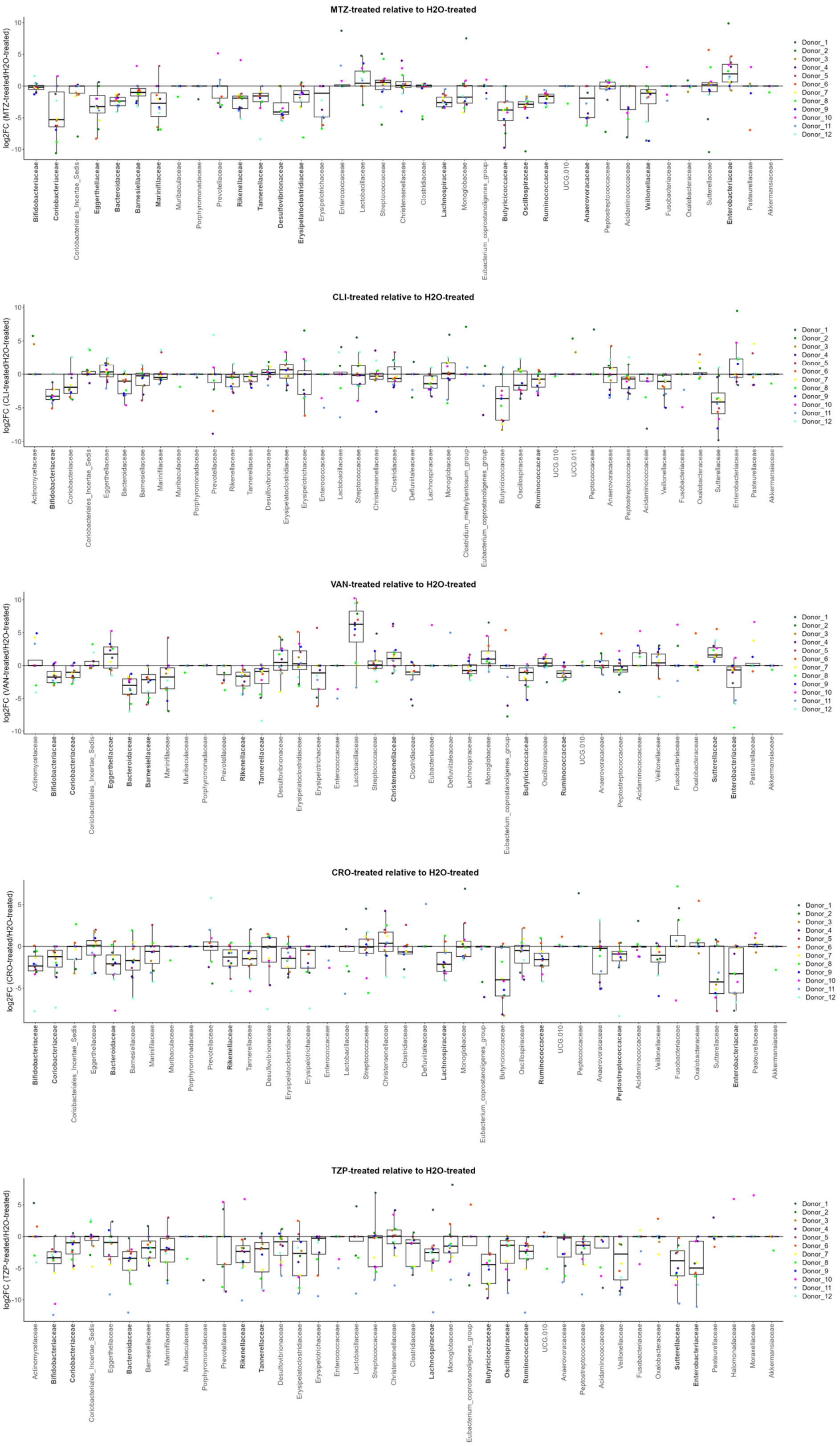


**<u>Figure S3</u>: Log2FC of bacterial families in antibiotic-treated faecal cultures as compared to water-treated faecal cultures by individual donor.** Bacterial families shown in bold font on the x-axis had a significant change in antibiotic-treated faecal cultures as compared to water-treated faecal cultures. n = 12 healthy human faecal donors, Wilcoxon signed rank test (two-sided) of log-transformed abundances with Benjamini Hochberg false discovery rate (FDR) correction, $p < 0.05$. MTZ = metronidazole, CLI = clindamycin, VAN = vancomycin, CRO = ceftriaxone, TZP = piperacillin/tazobactam.

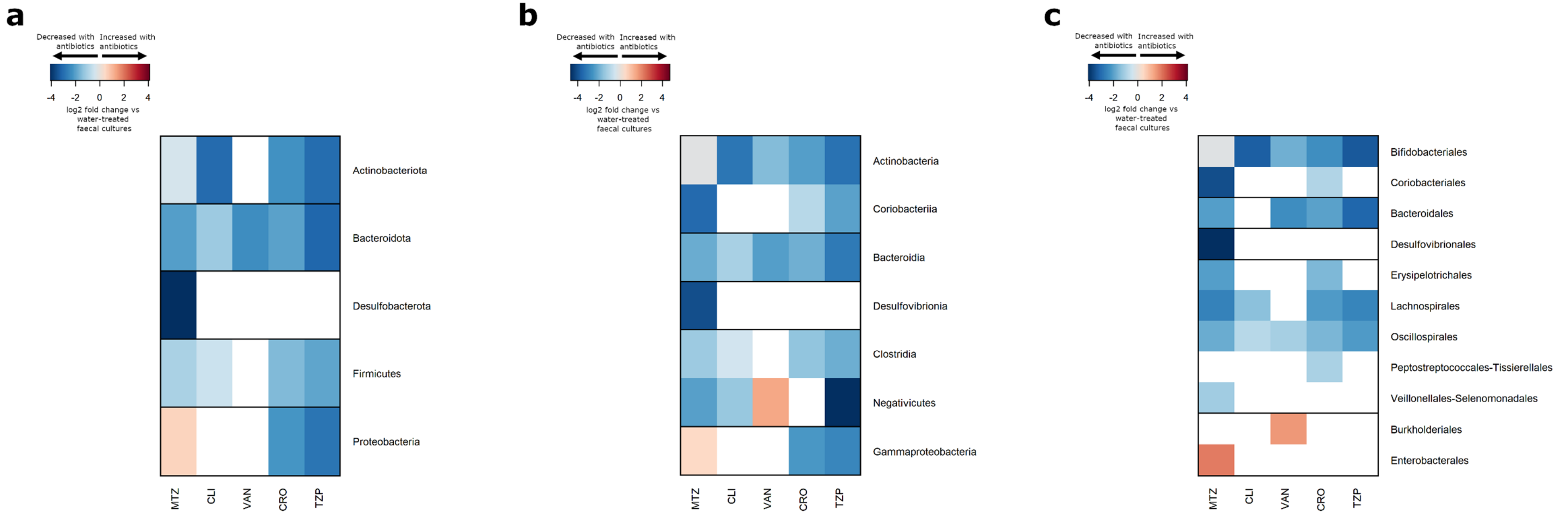


**<u>Figure S4</u>: Antibiotic treatment significantly reduced the abundance of gut commensals at the phylum, class, and order levels.** Heatmaps illustrating the log2 fold change (antibiotic-treated relative to water-treated) in bacterial taxa that were significantly decreased (shown in blue) or increased (shown in red) in faecal cultures following treatment with antibiotics that promote VRE intestinal colonisation. Taxa that were not significantly changed with antibiotic treatment were not plotted (shown in white). Changes are plotted at **(a)** phylum level, **(b)** class level, and **(c)** order level. n = 12 healthy human faecal donors, Wilcoxon signed rank test (two-sided) of log-transformed abundances with Benjamini Hochberg false discovery rate (FDR) correction, $p < 0.05$. MTZ = metronidazole, CLI = clindamycin, VAN = vancomycin, CRO = ceftriaxone, TZP = piperacillin/tazobactam.

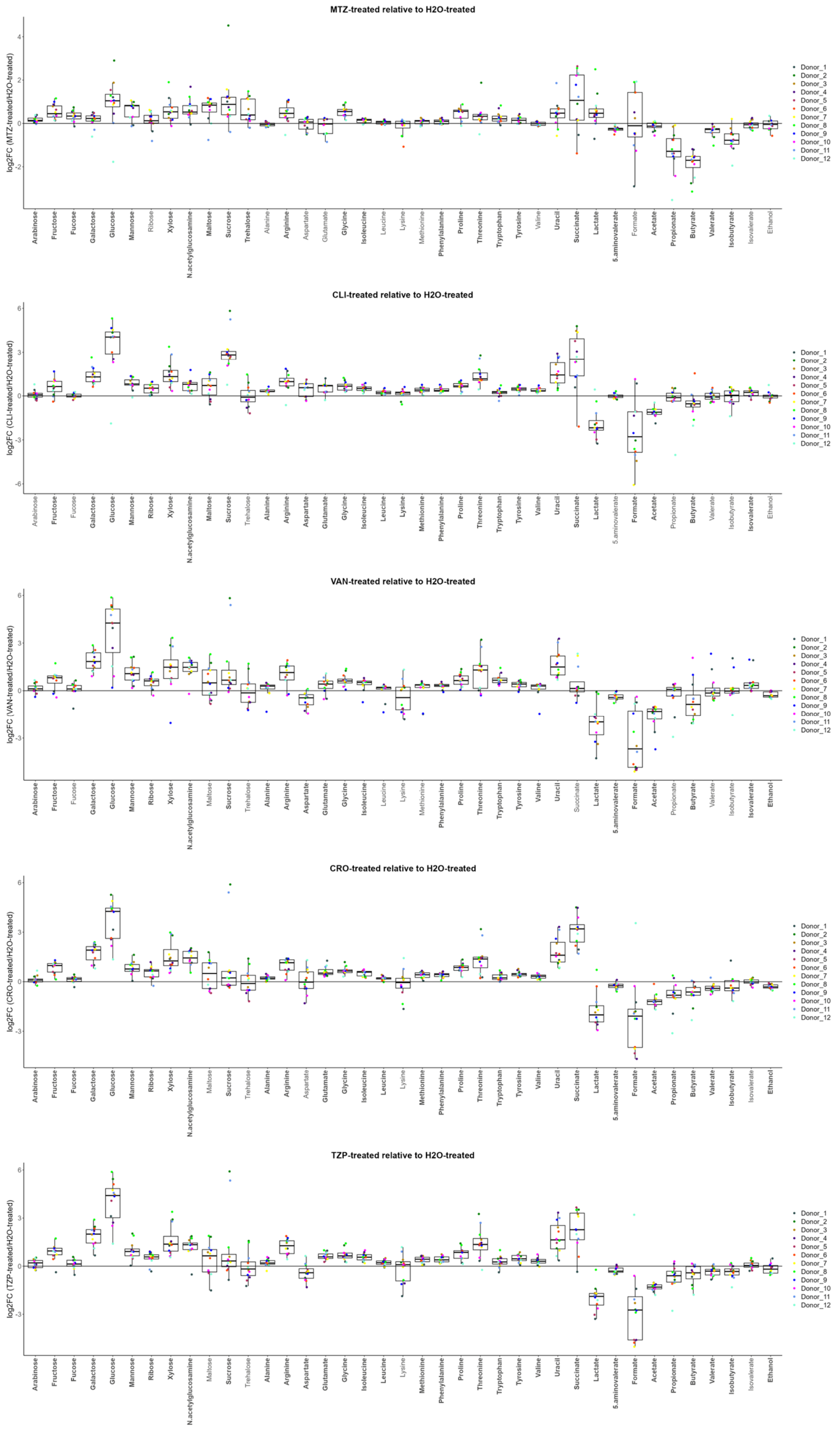


**Figure S5: Log2FC of nutrients and metabolites in antibiotic-treated faecal cultures as compared to water-treated faecal cultures by individual donor.** Nutrients or metabolites shown in bold font on the x-axis had a significant change in antibiotic-treated faecal cultures as compared to water-treated faecal cultures. n = 12 healthy human faecal donors, Wilcoxon signed rank test (two-sided) of log-transformed abundances with Benjamini Hochberg false discovery rate (FDR) correction, $p < 0.05$. MTZ = metronidazole, CLI = clindamycin, VAN = vancomycin, CRO = ceftriaxone, TZP = piperacillin/tazobactam.

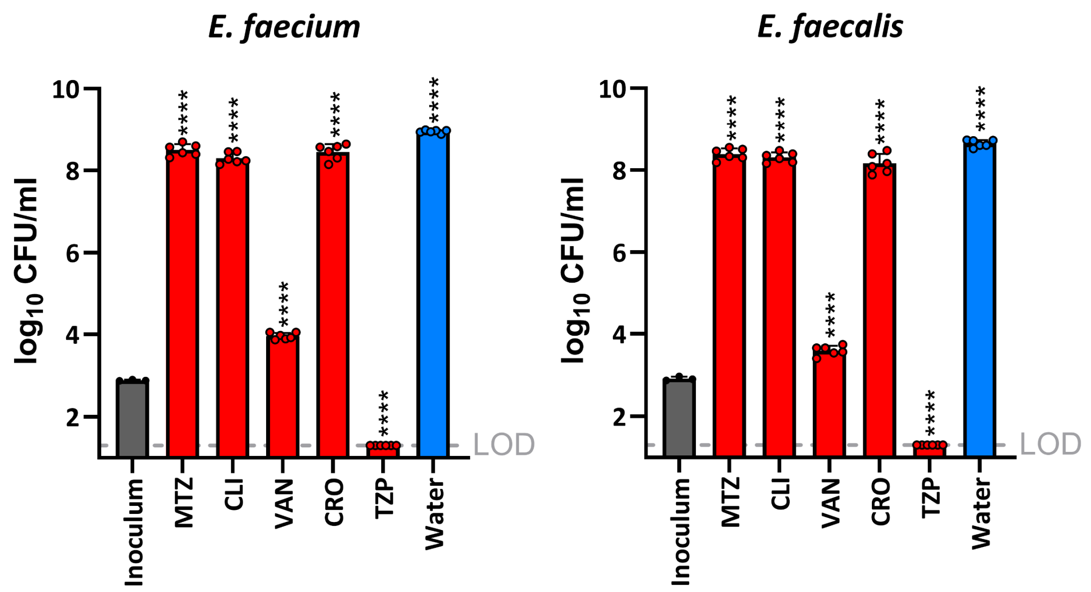


**<u>Figure S6</u>: VRE strains can grow in faecal concentrations of MTZ, CLI, VAN and CRO but are killed by faecal concentrations of TZP.** Vancomycin-resistant *E. faecium* (NCTC 12202) and *E. faecalis* (NCTC 12201) were grown anaerobically in gut growth medium supplemented with water or with one of the following antibiotics at concentrations mimicking those measured in human faeces: metronidazole (MTZ), clindamycin (CLI), vancomycin (VAN), ceftriaxone (CRO), or piperacillin/tazobactam (TZP). Antibiotic- or water-treated samples were compared to the inoculum using a one-way ANOVA with Dunnett's multiple comparison test. n=6 replicates from 3 independent experiments for the antibiotic- or water-treated groups, n=3 replicates from 3 independent experiments from the inoculum group, with data shown as mean ± SD. LOD = limit of detection.

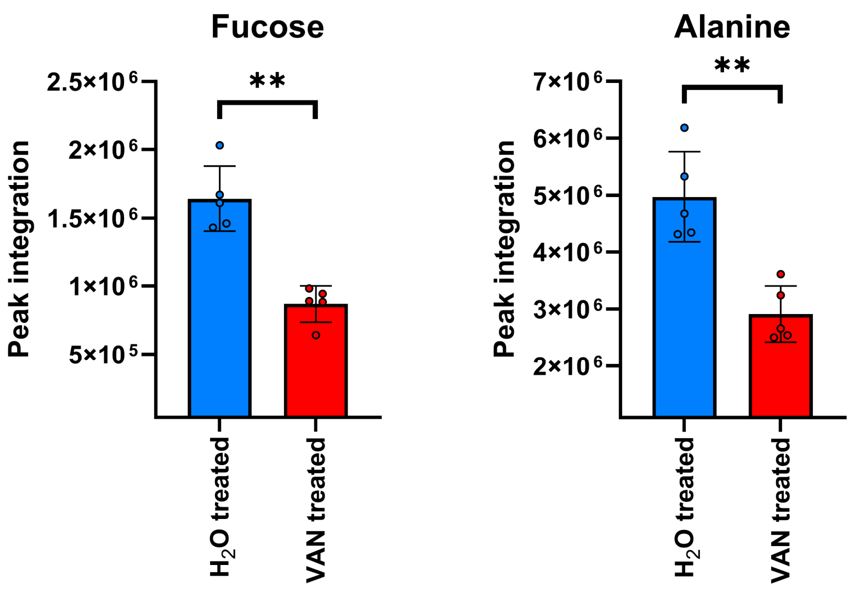


**<u>Figure S7</u>: Nutrients decreased in mouse faeces with vancomycin treatment.** Comparison of nutrients measured in VAN-treated mouse faeces to $H_2O$-treated mouse faeces used an unpaired t-test (two-sided) with Benjamini Hochberg FDR. n = 5 mice per group from one independent experiment, with data shown as mean ± SD. ** = P ≤ 0.01.

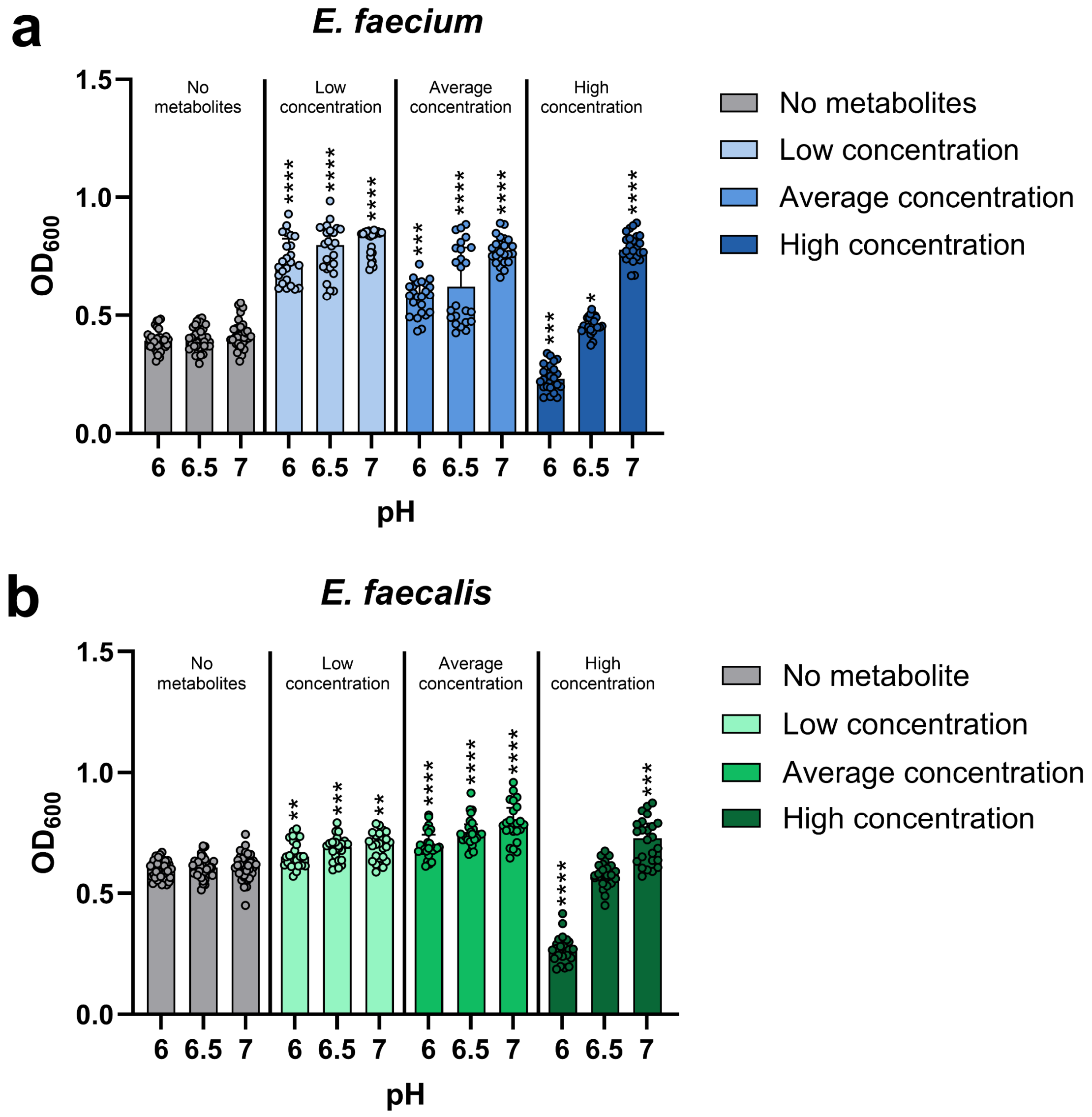


**Figure S8: VRE growth was suppressed at pH 6 by a mixture of 10 microbial metabolites at high concentrations.** Vancomycin-resistant *E. faecium* (NCTC 12202) and *E. faecalis* (NCTC 12201) were grown in tryptic soy broth supplemented with a mixture of 10 metabolites mimicking the low, average, or high concentrations measured in healthy human faeces, or unsupplemented ("No metabolite" control). Broth was adjusted to pH 6, 6.5, or 7 to mimic the pH of the healthy large intestine, and cultures were incubated under anaerobic conditions overnight. Data shown as medians ± IQR, with 24-36 replicates from 3 independent experiments. Kruskal-Wallis with Dunn's multiple comparison test comparing the no metabolite control to each concentration of the metabolite mixture at the same pH. * $P \leq 0.05$, ** = $P \leq 0.01$, *** = $P \leq 0.001$, **** = $P \leq 0.0001$. $OD_{600}$ = optical density at 600 nm.

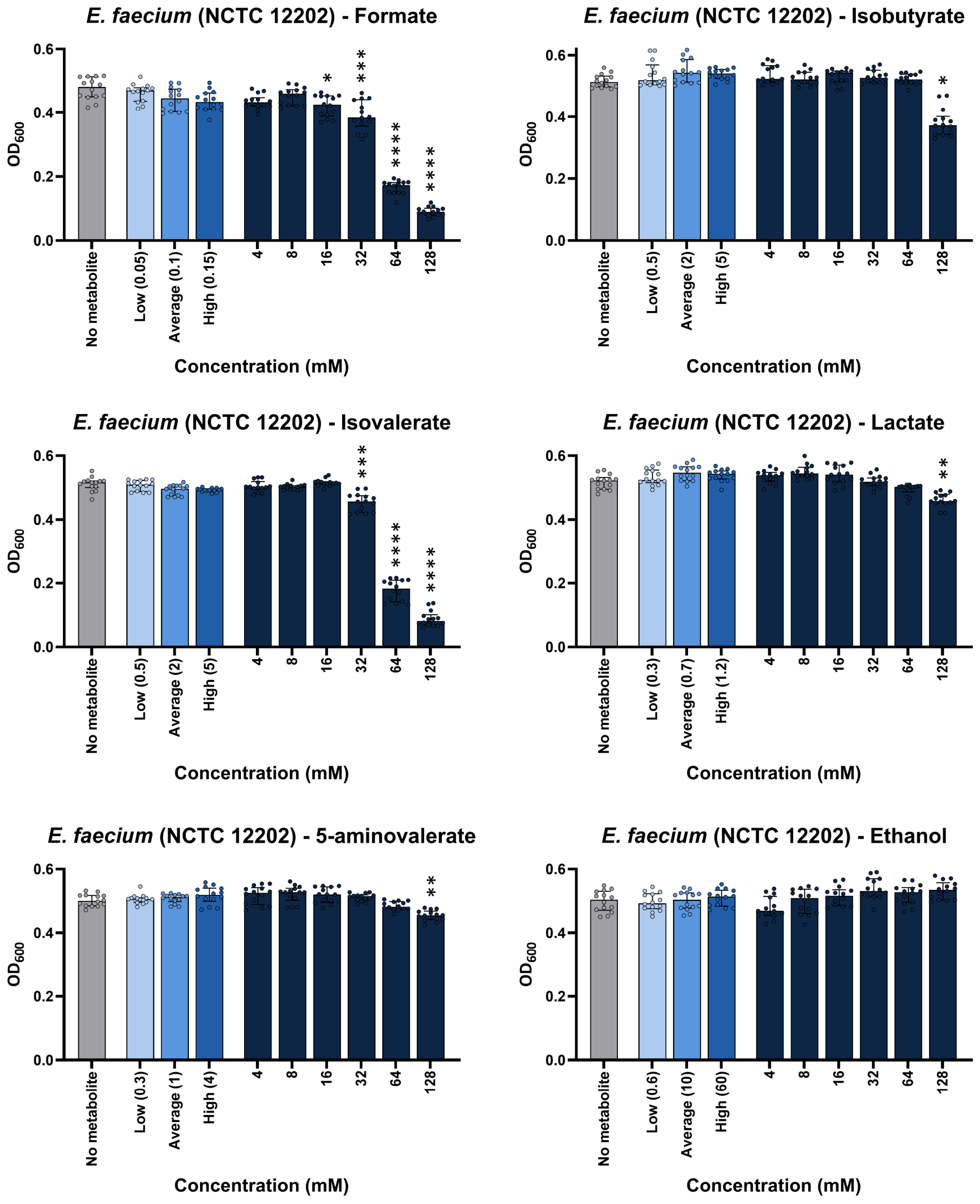


**Figure S9: Some individual microbial metabolites did not suppress vancomycin-resistant *E. faecium* (NCTC 12202) growth at physiological concentrations.** Vancomycin-resistant *E. faecium* (NCTC 12202) was grown in tryptic soy broth (pH 6) supplemented with an individual metabolite at low, average or high concentrations measured in healthy human faeces, concentrations spanning a defined range (4-128 mM), or unsupplemented (no metabolite control). Cultures were incubated under anaerobic conditions overnight. Growth was measured with 14 replicates in two independent experiments. Kruskal-Wallis test with Dunn's multiple comparison test (each metabolite was compared to the no metabolite control). * = $P \leq 0.05$, ** = $P \leq 0.01$, *** = $P \leq 0.001$, **** = $P \leq 0.0001$. Data are presented as medians ± IQR. Optical density at 600 nm, $OD_{600}$.

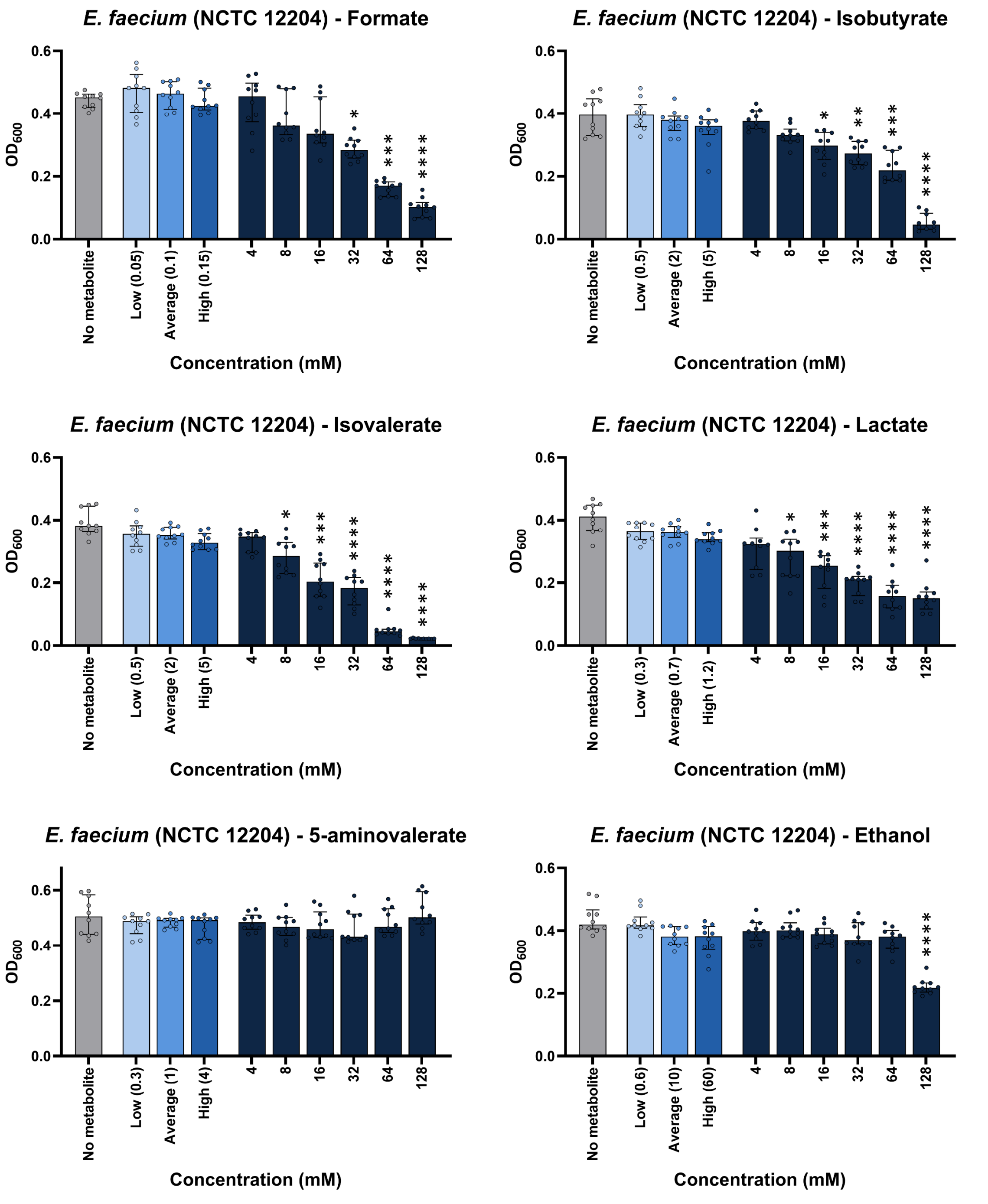


**Figure S10: Some individual microbial metabolites did not suppress vancomycin-resistant *E. faecium* (NCTC 12204) growth at physiological concentrations.** Vancomycin-resistant *E. faecium* (NCTC 12204) was grown in tryptic soy broth (pH 6) supplemented with an individual metabolite at low, average or high concentrations measured in healthy human faeces, concentrations spanning a defined range (4-128 mM), or unsupplemented (no metabolite control). Cultures were incubated under anaerobic conditions overnight. Growth was measured with 10 replicates in two independent experiments. Kruskal-Wallis test with Dunn's multiple comparison test (each metabolite was compared to the no metabolite control). * = $P \leq 0.05$, ** = $P \leq 0.01$, *** = $P \leq 0.001$, **** = $P \leq 0.0001$. Data are presented as medians ± IQR. Optical density at 600 nm, $OD_{600}$.

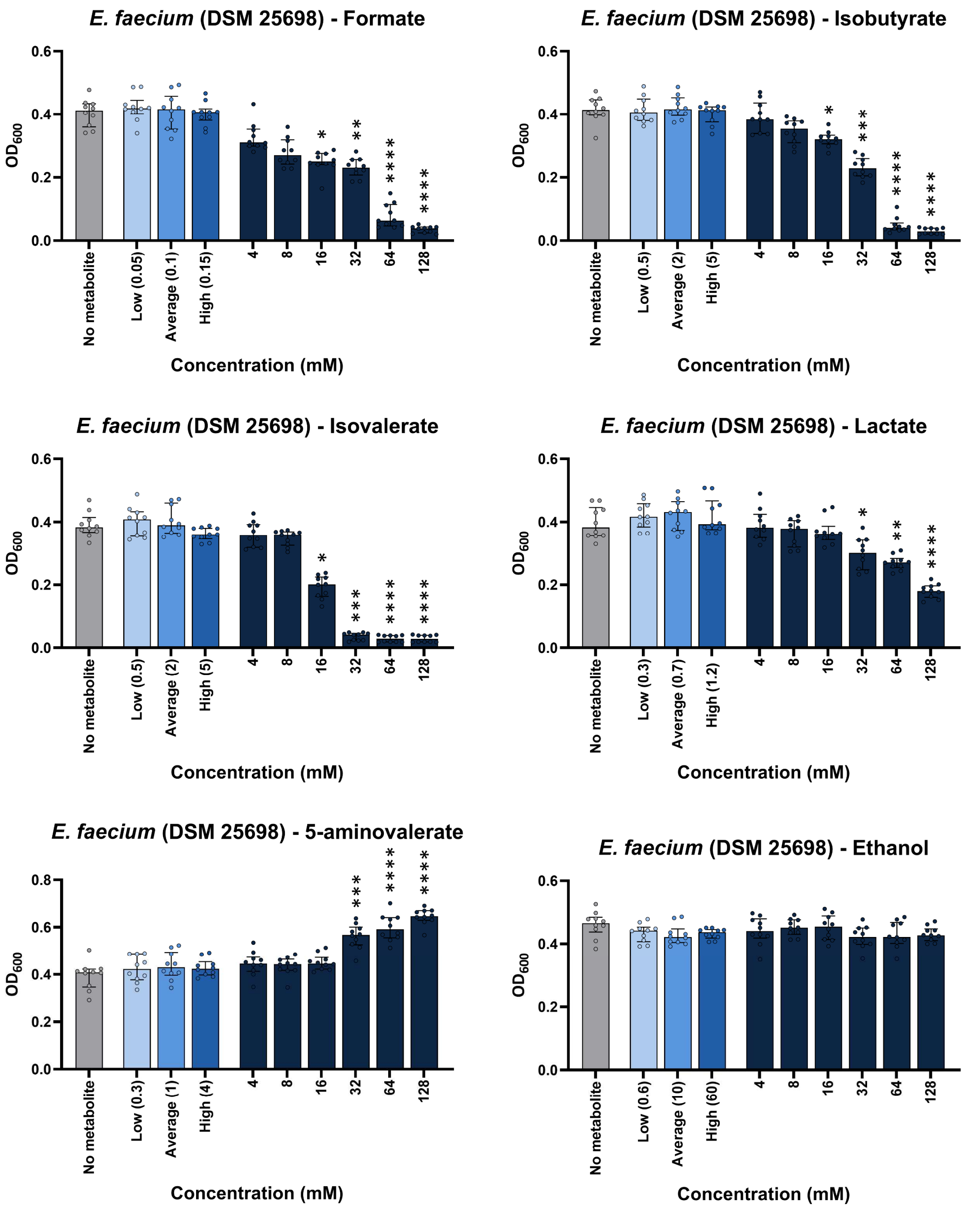


**<u>Figure S11</u>: Some individual microbial metabolites did not suppress vancomycin-resistant *E. faecium* (DSM 25698) growth at physiological concentrations.** Vancomycin-resistant *E. faecium* (DSM 25698) was grown in tryptic soy broth (pH 6) supplemented with an individual metabolite at low, average or high concentrations measured in healthy human faeces, concentrations spanning a defined range (4-128 mM), or unsupplemented (no metabolite control). Cultures were incubated under anaerobic conditions overnight. Growth was measured with 10 replicates in two independent experiments. Kruskal-Wallis test with Dunn's multiple comparison test (each metabolite was compared to the no metabolite control). * = $P \leq 0.05$, ** = $P \leq 0.01$, *** = $P \leq 0.001$, **** = $P \leq 0.0001$. Data are presented as medians ± IQR. Optical density at 600 nm, $OD_{600}$.

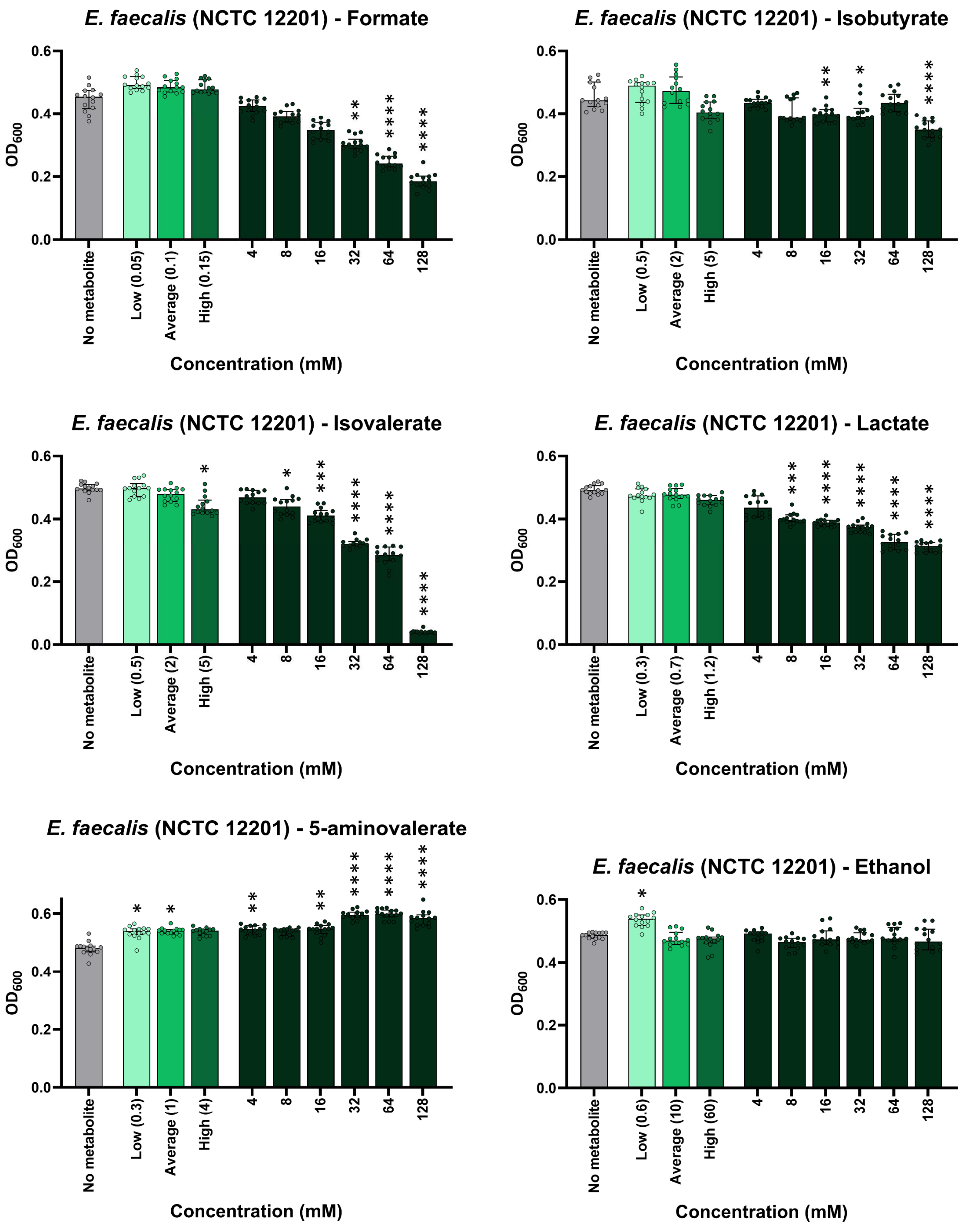


<u>Figure S12</u>**: Some individual microbial metabolites did not suppress vancomycin-resistant *E. faecalis* (NCTC 12201) growth at physiological concentrations.** Vancomycin-resistant *E. faecalis* (NCTC 12201) was grown in tryptic soy broth (pH 6) supplemented with an individual metabolite at low, average or high concentrations measured in healthy human faeces, concentrations spanning a defined range (4-128 mM), or unsupplemented (no metabolite control). Cultures were incubated under anaerobic conditions overnight. Growth was measured with 14 replicates in two independent experiments. Kruskal-Wallis test with Dunn's multiple comparison test (each metabolite was compared to the no metabolite control). * = $P \leq 0.05$, ** = $P \leq 0.01$, *** = $P \leq 0.001$, **** = $P \leq 0.0001$. Data are presented as medians ± IQR. Optical density at 600 nm, $OD_{600}$.

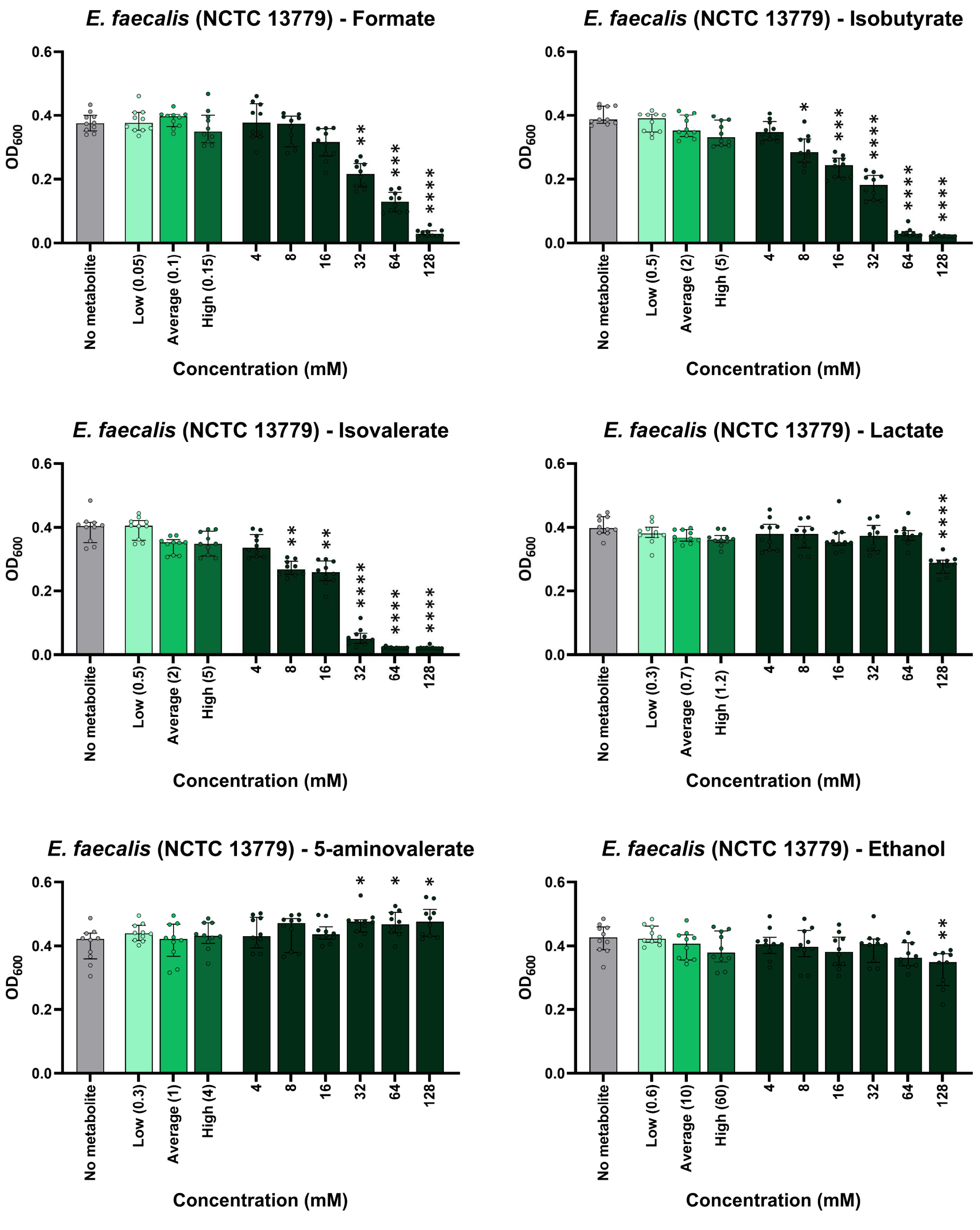


**<u>Figure S13</u>: Some individual microbial metabolites did not suppress vancomycin-resistant *E. faecalis* (NCTC 13779) growth at physiological concentrations.** Vancomycin-resistant *E. faecalis* (NCTC 13779) was grown in tryptic soy broth (pH 6) supplemented with an individual metabolite at low, average or high concentrations measured in healthy human faeces, concentrations spanning a defined range (4-128 mM), or unsupplemented (no metabolite control). Cultures were incubated under anaerobic conditions overnight. Growth was measured with 10 replicates in two independent experiments. Kruskal-Wallis test with Dunn's multiple comparison test (each metabolite was compared to the no metabolite control). * = $P \leq 0.05$, ** = $P \leq 0.01$, *** = $P \leq 0.001$, **** = $P \leq 0.0001$. Data are presented as medians ± IQR. Optical density at 600 nm, $OD_{600}$.

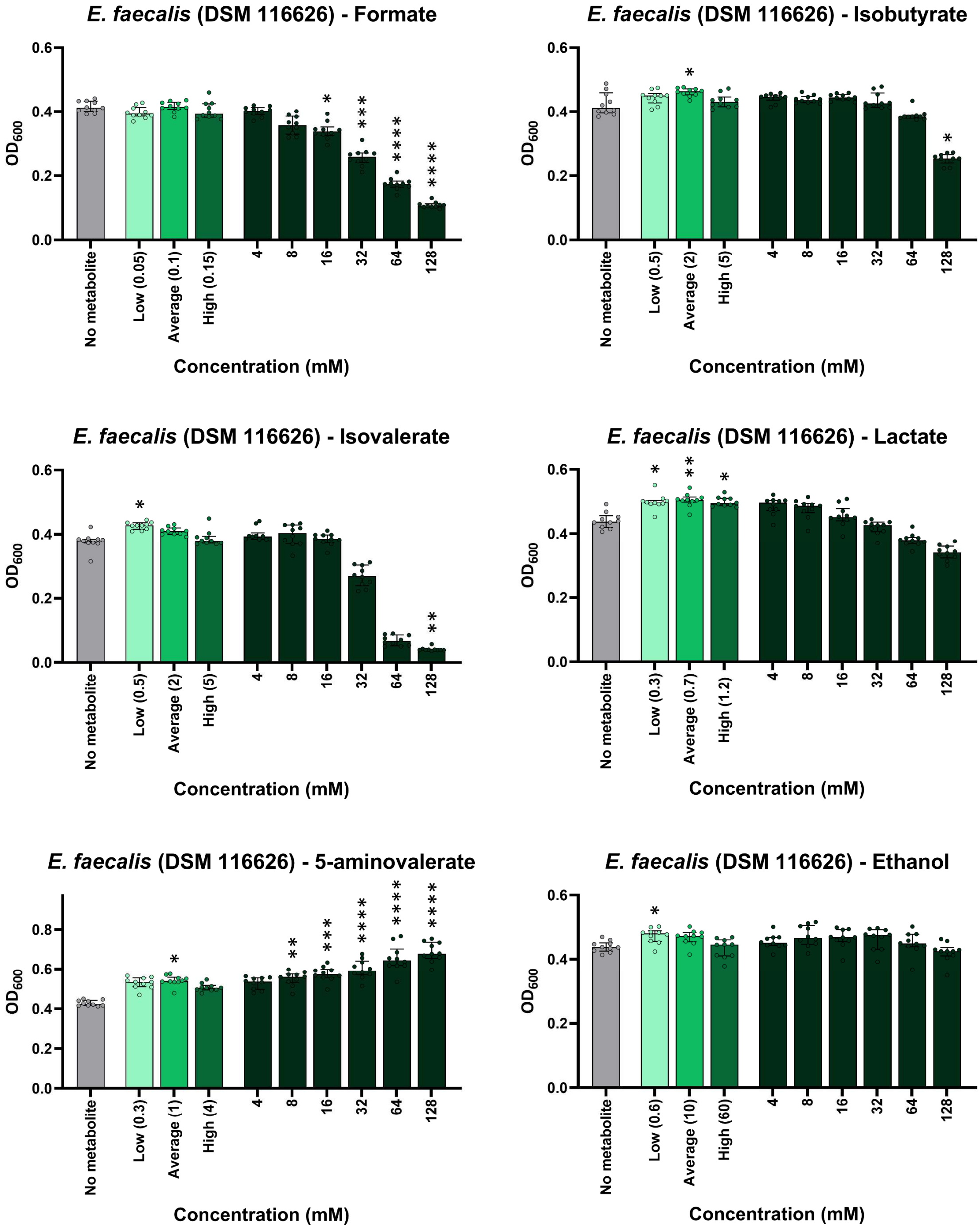


**<u>Figure S14</u>: Some individual microbial metabolites did not suppress vancomycin-resistant *E. faecalis* (DSM 116626) growth at physiological concentrations.** Vancomycin-resistant *E. faecalis* (DSM 116626) was grown in tryptic soy broth (pH 6) supplemented with an individual metabolite at low, average or high concentrations measured in healthy human faeces, concentrations spanning a defined range (4-128 mM), or unsupplemented (no metabolite control). Cultures were incubated under anaerobic conditions overnight. Growth was measured with 10 replicates in two independent experiments. Kruskal-Wallis test with Dunn's multiple comparison test (each metabolite was compared to the no metabolite control). * = $P \leq 0.05$, ** = $P \leq 0.01$, *** = $P \leq 0.001$, **** = $P \leq 0.0001$. Data are presented as medians ± IQR. Optical density at 600 nm, $OD_{600}$.

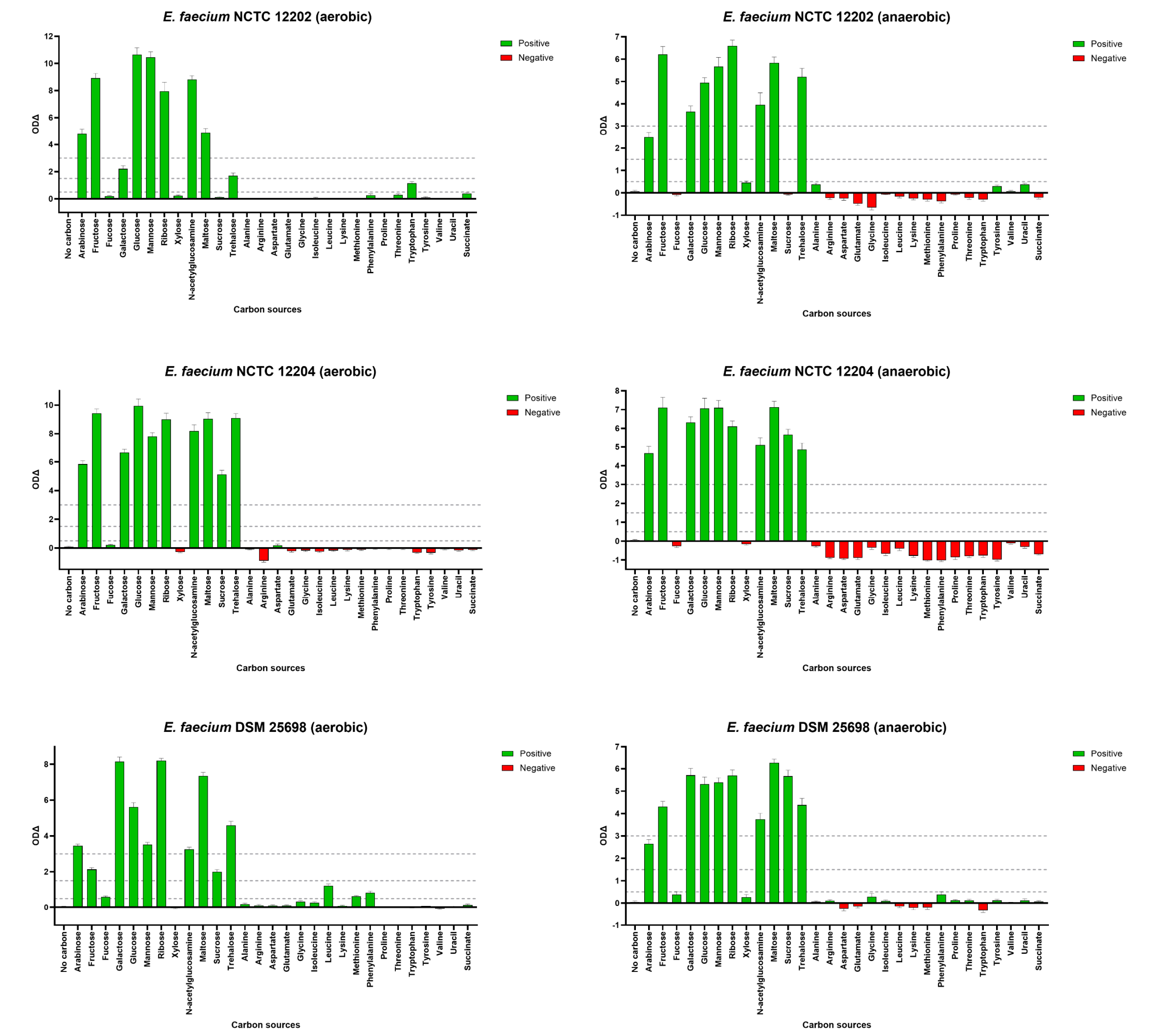


<u>Figure S15</u>: **Vancomycin-resistant *E. faecium* growth was supported by individual nutrients found in antibiotic-treated faecal microbiomes as carbon sources.** Functional differences (ODΔ) between growth of vancomycin-resistant *E. faecium* on individual carbon sources under both aerobic and anaerobic conditions. ODΔ shows the sum of functional differences between the individual carbon source growth curve compared to growth on the negative (no carbon) control. High growth is indicated by an ODΔ > 3, moderate growth is indicated by an ODΔ between 1.5 and 3, low growth is indicated by an ODΔ between 0.5 and 1.5, and negligible growth or no growth is indicated by an ODΔ < 0.5. n = 6 replicates in 2-3 independent experiments. The error bars represent the 95% confidence intervals.

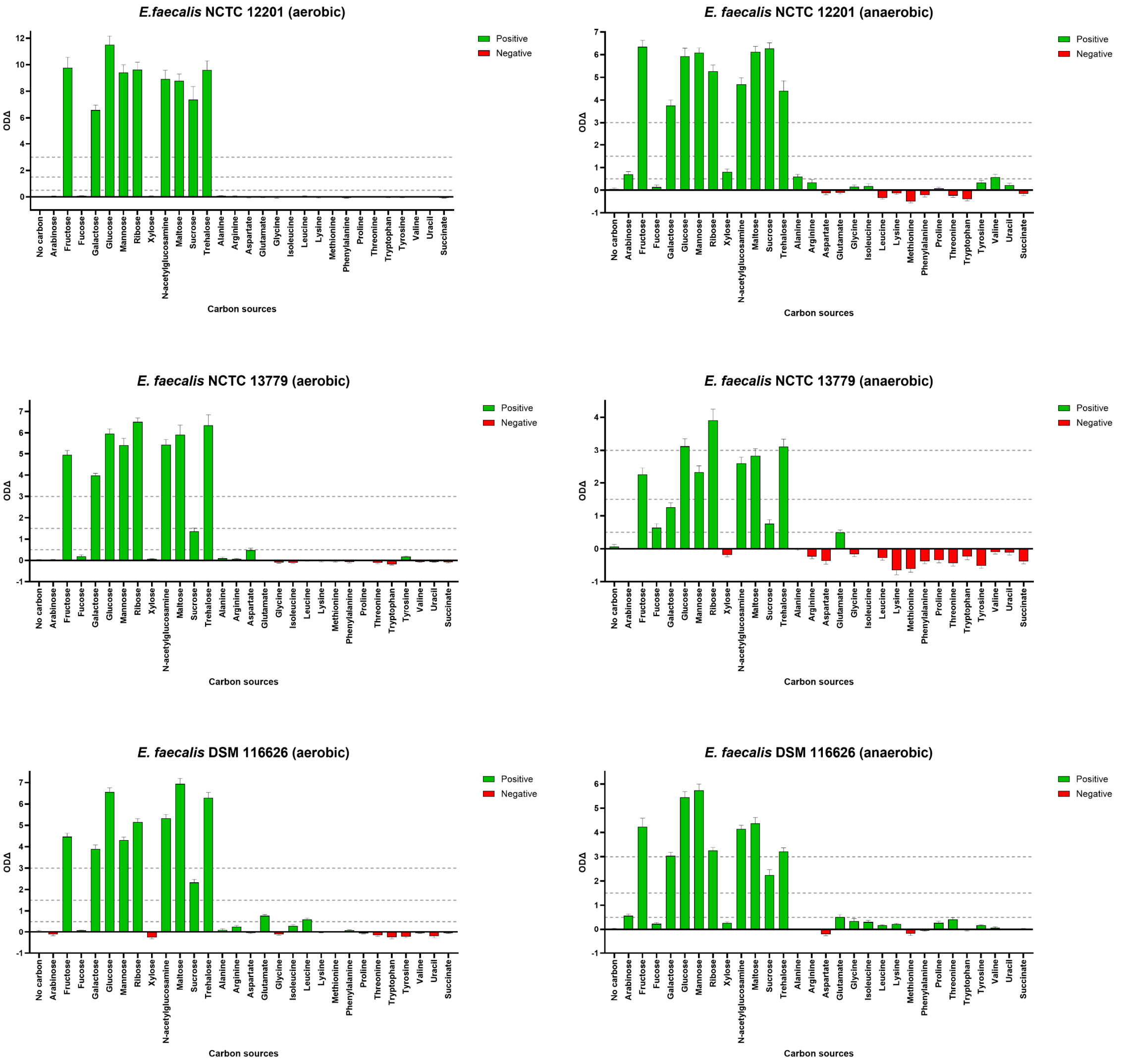


**<u>Figure S16</u>: Vancomycin-resistant *E. faecalis* growth was supported by individual nutrients found in antibiotic-treated faecal microbiomes as carbon sources.** Functional differences (ODΔ) between growth of vancomycin-resistant *E. faecalis* on individual carbon sources under both aerobic and anaerobic conditions. ODΔ shows the sum of functional differences between the individual carbon source growth curve compared to growth on the negative (no carbon) control. High growth is indicated by an ODΔ > 3, moderate growth is indicated by an ODΔ between 1.5 and 3, low growth is indicated by an ODΔ between 0.5 and 1.5, and negligible growth or no growth is indicated by an ODΔ < 0.5. n = 6 replicates in 2-3 independent experiments. The error bars represent the 95% confidence intervals.

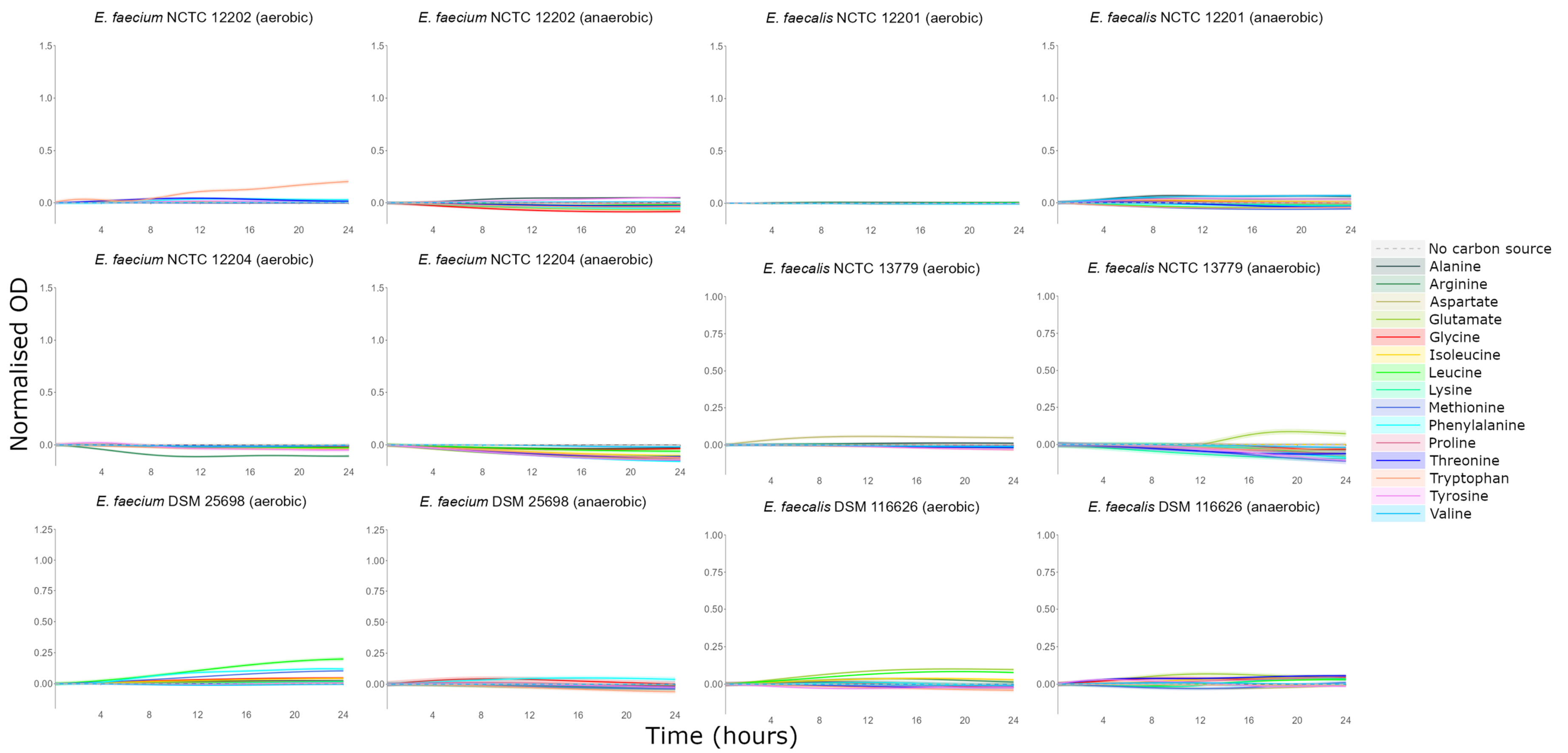


**Figure S17: Amino acids supported low levels of VRE growth or did not support VRE growth as carbon sources.** AMiGA-predicted growth curves for vancomycin-resistant *E. faecium* and vancomycin-resistant *E. faecalis* strains cultured in minimal medium supplemented with single carbon sources under anaerobic or aerobic conditions. The bold lines show the predicted mean of growth while the shaded bands show the predicted 95% credible intervals. n = 6 replicates in 2–3 independent experiments. OD = optical density.

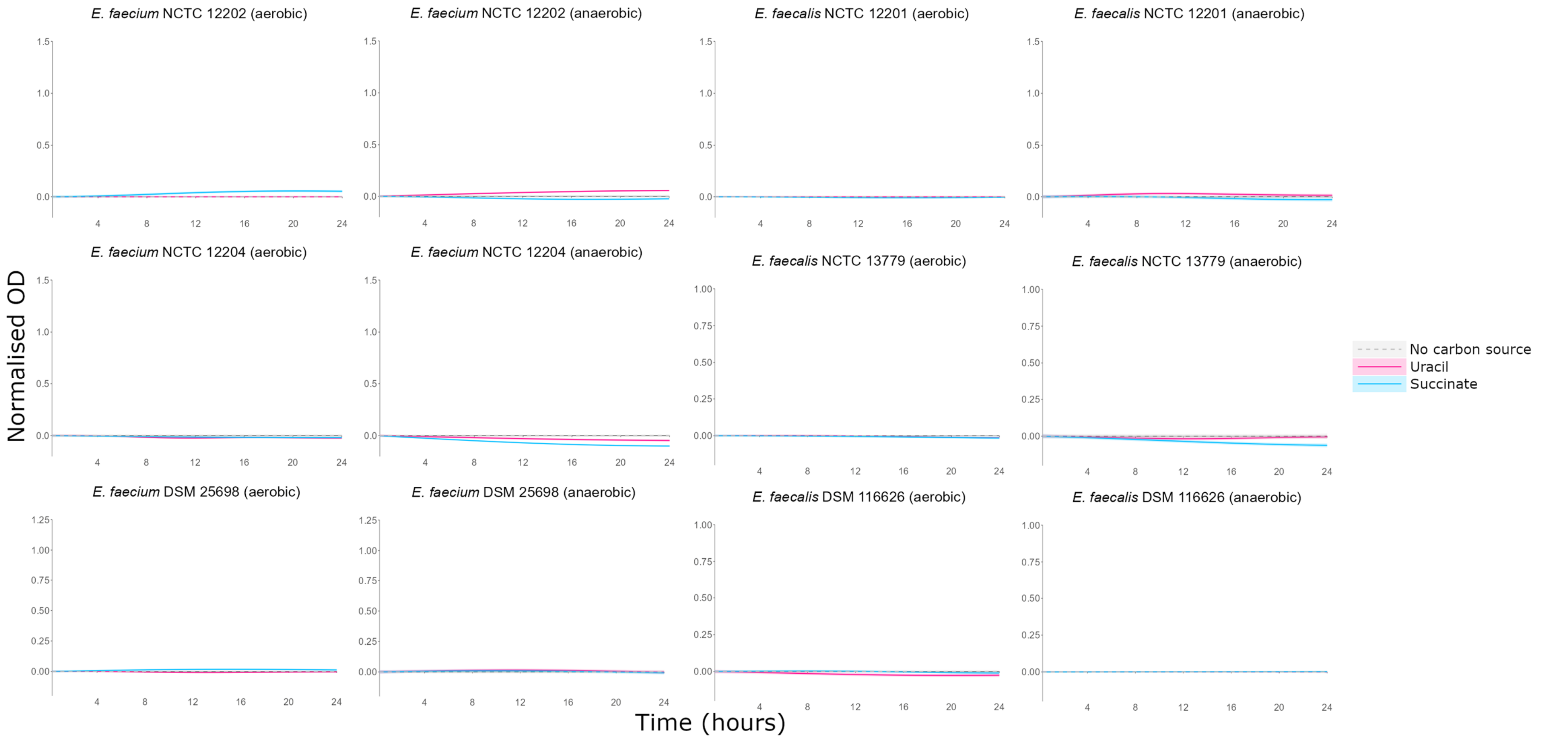


**<u>Figure S18</u>: Uracil and succinate did not support VRE growth as carbon sources.** AMiGA-predicted growth curves for vancomycin-resistant *E. faecium* and vancomycin-resistant *E. faecalis* strains cultured in minimal medium supplemented with single carbon sources under anaerobic or aerobic conditions. The bold lines show the predicted mean of growth while the shaded bands show the predicted 95% credible intervals. n = 6 replicates in 2–3 independent experiments. OD = optical density.

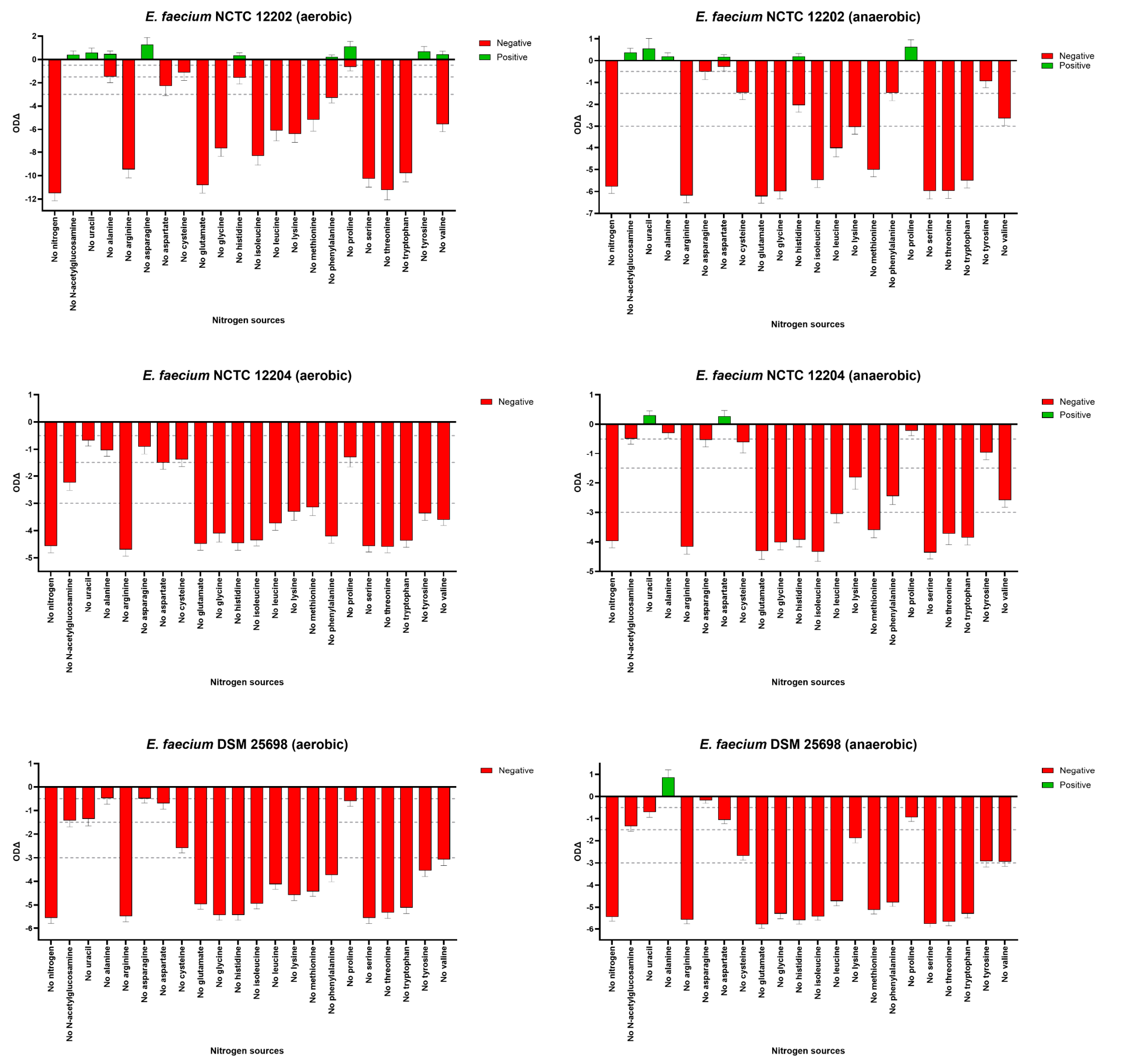


**<u>Figure S19</u>: Vancomycin-resistant *E. faecium* growth was supported by individual nutrients found in antibiotic-treated faecal microbiomes as nitrogen sources.** Functional differences (ODΔ) between growth of vancomycin-resistant *E. faecium* on nitrogen sources in a leave-one-out experiment under both aerobic and anaerobic conditions. ODΔ shows the sum of functional differences between the growth curve for each leave-one-out mixture (lacking a single nitrogen source) compared to the growth curve for the full mixture of nitrogen sources. A high decrease in growth is indicated by an ODΔ < -3, a moderate decrease in growth is indicated by an ODΔ between -1.5 and -3, a low decrease in growth is indicated by an ODΔ between -0.5 and -1.5, and negligible or no decrease in growth is indicated by an ODΔ > -0.5. n = 6 replicates in 2-3 independent experiments. The error bars represent the 95% confidence intervals.

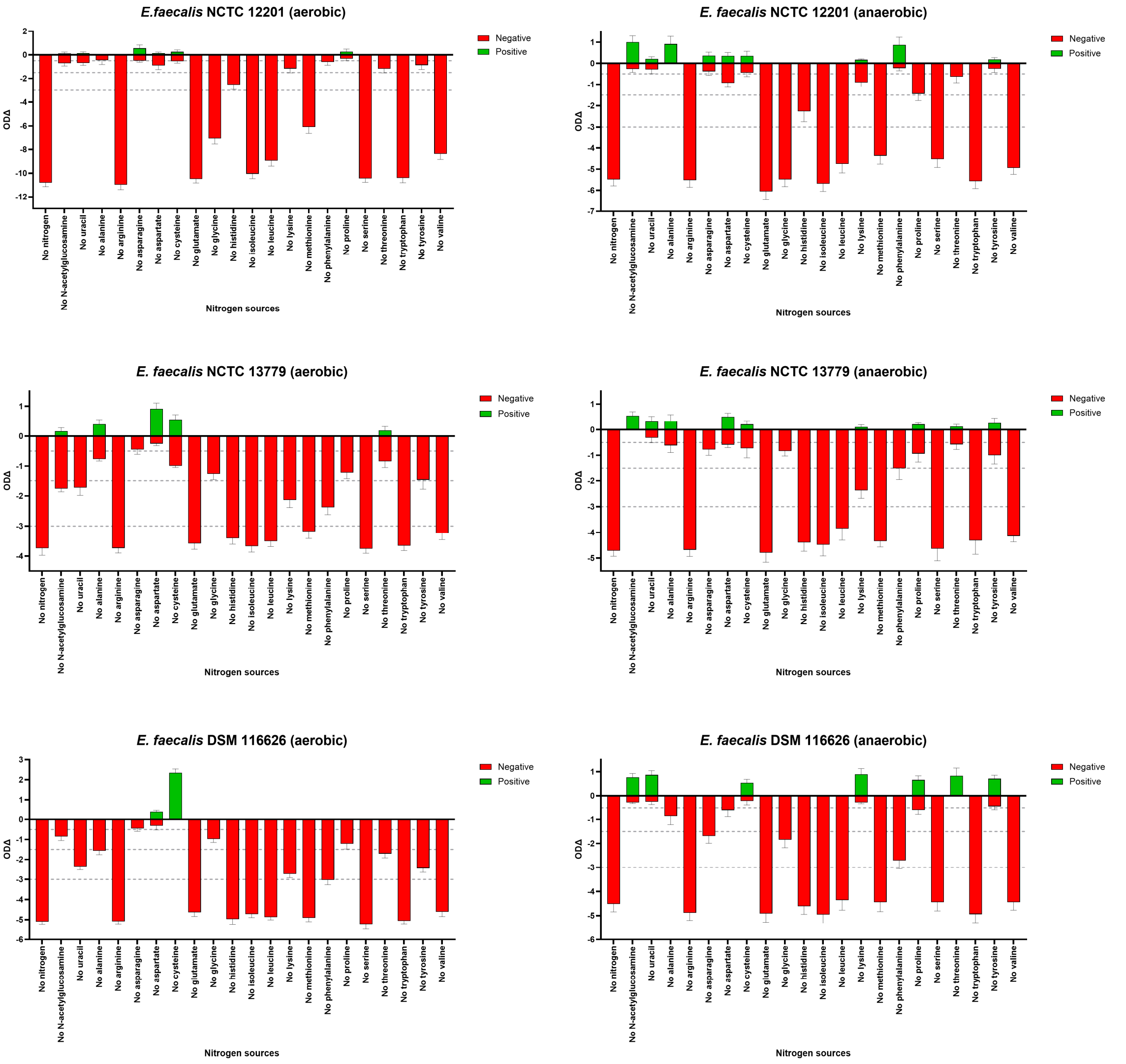


**<u>Figure S20</u>: Vancomycin-resistant *E. faecalis* growth was supported by individual nutrients found in antibiotic-treated faecal microbiomes as nitrogen sources.** Functional differences (ODΔ) between growth of vancomycin-resistant *E. faecalis* on nitrogen sources in a leave-one-out experiment under both aerobic and anaerobic conditions. ODΔ shows the sum of functional differences between the growth curve for each leave-one-out mixture (lacking a single nitrogen source) compared to the growth curve for the full mixture of nitrogen sources. A high decrease in growth is indicated by an ODΔ < -3, a moderate decrease in growth is indicated by an ODΔ between -1.5 and -3, a low decrease in growth is indicated by an ODΔ between -0.5 and -1.5, and negligible or no decrease in growth is indicated by an ODΔ > -0.5. n = 6 replicates in 2-3 independent experiments. The error bars represent the 95% confidence intervals.

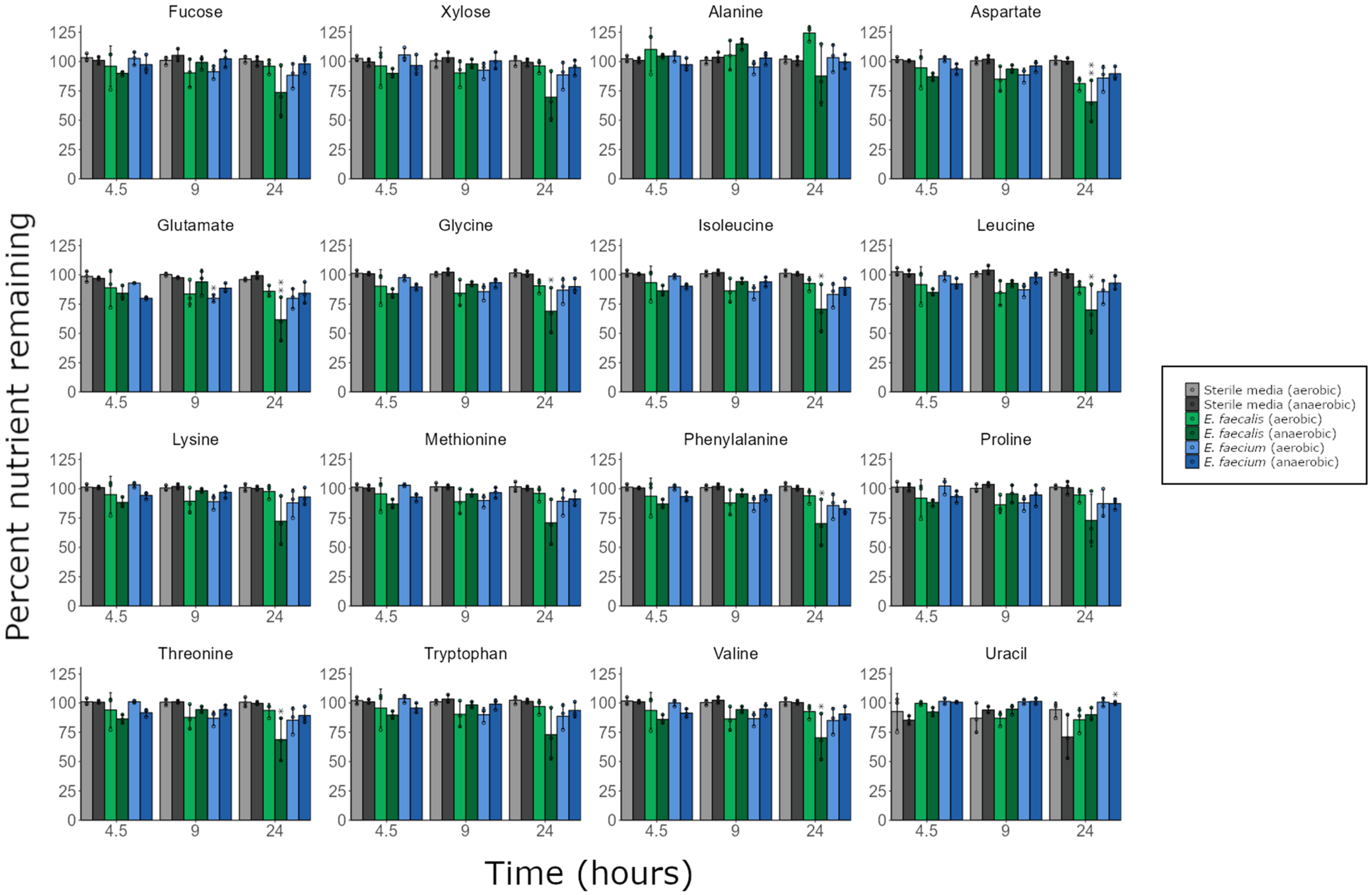


**Figure S21: VRE did not prefer some nutrients over others when provided with a mixture of nutrients.** Percentage of nutrients remaining in vancomycin-resistant *E. faecium* (NCTC 12202) and *E. faecalis* (NCTC 12201) cultures grown in a minimal medium supplemented with a mixture of nutrients, under aerobic or anaerobic conditions. $^{1}$H-NMR spectroscopy peak integrations were used to measure nutrient concentrations. Comparisons were made using a two-way mixed ANOVA followed by pairwise comparisons with Bonferroni correction. n = 3 replicates per group, with data shown as mean ± SD. * = $P \leq 0.05$, ** = $P \leq 0.01$.
